# Single-Shot Design of a Cyclic Peptide Inhibitor of HIV-1 Membrane Fusion with EvoBind

**DOI:** 10.1101/2025.04.30.651413

**Authors:** Diandra Daumiller, Federica Giammarino, Qiuzhen Li, Anders Sönnerborg, Rafael Ceña Diez, Patrick Bryant

## Abstract

HIV evades the immune system through rapid mutation of its surface proteins, particularly the envelope glycoprotein. However, the core mechanism of viral entry, CD4 binding and co-receptor engagement remains conserved. While therapies such as Lenacapavir represent important advances, the continued emergence of resistant strains will demand new and more adaptable treatment strategies. This challenge is not unique to HIV; future pandemics will likely present similar pressures, highlighting the need for drug design methods that are not only effective but also fast and scalable. Recent advances in protein structure prediction have transformed the landscape of therapeutic design, enabling the accurate modelling of target structures from sequence alone and now facilitating the development of novel therapeutics without prior structural data. EvoBind leverages these advances to rapidly generate cyclic peptide binders in a single design round, using only the amino acid sequence of a target protein. Cyclic peptides offer several advantages over traditional linear protein molecules, including increased stability, while their small size improves oral bioavailability and enables access to challenging binding sites. Here, we demonstrate the use of EvoBind to generate cyclic peptide binders against the HIV envelope protein gp41, which is essential for viral-host membrane fusion. Cell-based assays confirm potent inhibition of two different HIV-1 strains with no detectable toxicity. The combination of artificial intelligence-guided design and streamlined experimental validation can significantly accelerate therapeutic development, reduce costs, and provide timely solutions to the challenges posed by viral evolution and emerging global health threats.

## Introduction

The human immunodeficiency virus (HIV) remains a major global health challenge, with millions of people affected despite advances in antiviral therapy [1–3]. One of the primary difficulties in treating HIV is its ability to mutate rapidly, particularly in its envelope glycoproteins, which are key targets for neutralising antibodies [4–6]. However, some viral components, such as the capsid protein, remain structurally conserved and have been explored as alternative therapeutic targets [7,8]. Lenacapavir, a small-molecule inhibitor targeting HIV capsid assembly and disassembly, represents a breakthrough in long-acting antiretroviral therapy by disrupting viral replication at multiple stages [9,10].

While Lenacapavir provides an effective new strategy, new resistant strains will likely emerge, underscoring the need for continuously evolving therapeutic approaches [11]. Resistance selection experiments have identified several major drug resistance mutations in the HIV capsid (L56I, M66I, Q67H/Y, K70N/R, N74D/S, T107N) that reduce susceptibility to Lenacapavir. Except for Q67H, all these mutations reduced viral fitness. Most of these resistance mutations (M66I, Q67H, K70N/R, N74D/S) have also been observed in vivo in clinical trials [11–13].

Protein function is strictly dependent on tertiary and quaternary protein structure. Predicting protein structure from sequence has long been a major challenge, but the innovation introduced by AlphaFold has revolutionised the field [14] and laid the foundation for further advancements. By integrating AlphaFold with other machine learning models, it is now possible to predict additional protein properties, including different conformations [15,16], protein complexes [17,18], protein-ligand interactions [19–22], and protein binders [23–25].

Building on the foundation, new artificial intelligence tools are now being developed to not only predict structures but to design novel therapeutics. Even though we seem to be far away from accurately predicting small-molecule protein interactions without prior interface knowledge [26,27], protein binders can now be accurately designed and serve as alternatives. Particularly peptide-based drugs, which are known for their high specificity, appear promising [28,29]. Among these, cyclic peptides stand out due to their increased conformational stability, resistance to proteolytic degradation, and improved bioavailability compared to their linear counterparts [30,31].

To prepare for future pandemics and rapidly evolving pathogens, therapeutic development must be fast, accurate, and adaptable. Traditional drug discovery pipelines are not equipped to meet these demands [32]. In contrast, AI-based approaches allow for rapid exploration of therapeutic space, including the concept of *single-shot design*, where a potent therapeutic binder can be generated computationally in a single iteration without the need for extensive experimental screening or optimisation. In this work, we demonstrate the feasibility of this approach by applying EvoBind [23,32] to the de novo design of cyclic peptide binders targeting the HIV-1 envelope protein gp41 [33]. Given the well-documented issue of rapid resistance development in HIV [34], this approach represents a promising avenue for generating robust peptide-based therapeutics that can be swiftly adapted in response to emerging viral strains.

## Results

### Design of a Cyclic Peptide HIV-1 Inhibitor with EvoBind

Peptide, antibody and small molecule inhibitors are widely used to neutralise HIV, but the virus’s rapid evolution often enables it to escape from these therapeutic agents. Gp41, a transmembrane protein in the HIV envelope, plays a central role in viral fusion and entry by mediating the interaction between the virus and host cell membranes [35,36]. Using only the amino acid sequence of gp41 as input, without any prior specification of a target interface, we applied the AI-based framework EvoBind [23] to design cyclic peptide binders targeting this conserved viral component (Figure 1a).

**Figure 1.**
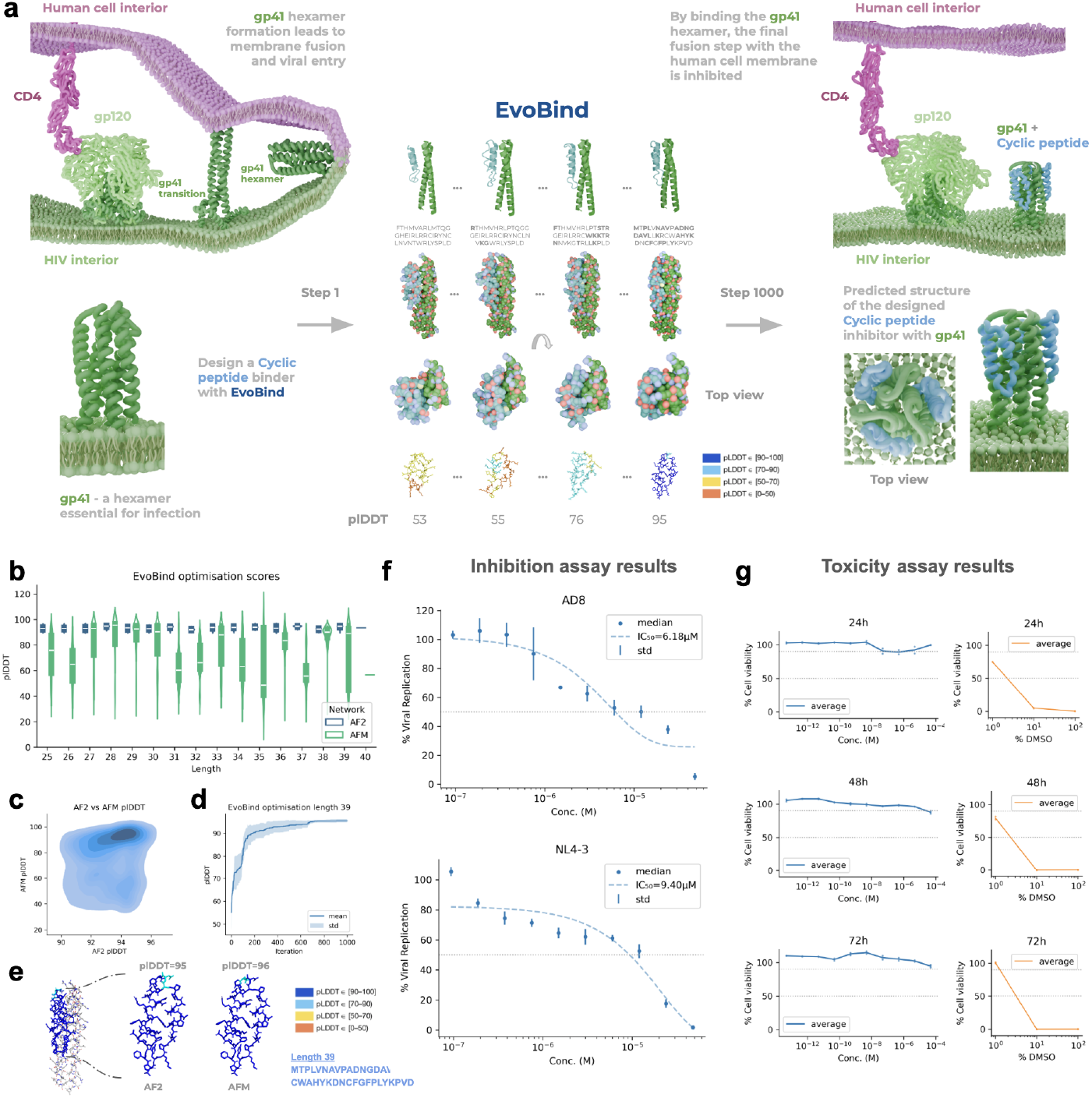
Design and inhibition mechanism of a cyclic peptide targeting HIV-1 fusion. **a)** Schematic overview of HIV-1 entry, highlighting the role of gp41 in membrane fusion, after CD4 (AF-P01730-F1-v4) and gp120 (PDB ID 8F7T) binding.The gp41 fusion hexamer (PDB ID 3CP1) is exposed only during a narrow temporal window, exactly when the human and HIV membranes are about to fuse. Starting from two out of six gp41 parts, constituting a “pseudo dimer” sequence, we use EvoBind to generate a cyclic peptide binder by iteratively improving predicted structure quality (plDDT) and interface compaction. Uniquely, EvoBind generates both the sequence and structure of the cyclic peptide in complex with the target simultaneously. By step 807, nearly all amino acids have been substituted from the random starting sequence, resulting in a stable, high-confidence binder conformation with a plDDT score of 95 compared to only 53 at step 1. The cyclic peptide is predicted to bind in triplicate to the gp41 hexamer, each engaging one pseudo-dimer and ultimately inhibiting the membrane fusion. **b)** Distribution of predicted confidence (plDDT) scores for designs with plDDT above 90 predicted with AlphaFold2 (AF2, blue) and comparison AlphaFold-multimer (AFM, green). The number of designs per length with AF2 plDDT above 90 is as follows: 25: 787, 26: 523, 27: 743, 28: 289, 29: 852, 30: 819, 31: 319, 32: 462, 33: 515, 34: 325, 35: 28, 36: 587, 37: 173, 38: 261, 39: 763, 40: 1. The white dots mark the medians and the box marks the quartiles with the lines extending to the remainder of the distribution. Agreement with AFM predictions varies substantially across lengths. All lengths yielded high plDDT scores for both AF2 and AFM network weights, except for length 40, which produced only one high-confidence design. Length 39 (L39) was selected as the final binder. **c)** Density plot (darker shade means more points) of AF2 vs AFM plDDT for protein-peptide complex prediction using all scores from Figure 1b (n=7447). The adapted prediction models using AF2 and AFM yield consistent plDDT scores across peptide lengths, supporting the reliability of the designed complexes. **d)** Highest plDDT so far vs iteration for the design of the evaluated peptide (length 39). The average across five different initialisations is displayed as a line with shaded standard deviation. **e)** Comparison between AF2 and AFM plDDT for the predictions of the design of length 39. The predicted protein-peptide complex structures show remarkable agreement between AF2 and AFM models. The receptor is shown in grey, and the peptide is colored by plDDT. AFM yields slightly higher plDDT scores overall (96 compared to 95 for AF2), indicating strong and consistent structural confidence. **f)** Inhibition data with displayed median and standard deviation at each measured concentration, based on n=4 measurements per concentration. The observed inhibition indicates successful target engagement under the constrained conditions of hexamer gp41 targeting, with IC_50_ values of 6.18 μM and 9.40 μM for the AD8 and NL4-3 HIV-1 strains, respectively (dotted grey line). **g)** Toxicity data at timepoints 24, 48 and 72 h, where the points represent each measurement (n=2 per concentration) and the line represents the average. Importantly, no cellular toxicity was detected even at high peptide concentrations, with cell viability remaining high 72 hours after exposure. In contrast, DMSO treatment resulted in complete cell death at >10%, confirming the sensitivity of the assay. These findings demonstrate that the designed cyclic peptide acts selectively on the virus without adversely affecting host cells. Cell viability is marked at 50 and 90% with dotted grey lines.

EvoBind begins from a randomly generated peptide sequence and iteratively introduces mutations to improve interface compactness and predicted confidence, measured by the predicted local distance difference test (plDDT [14]). Across 1000 rounds of mutation, the peptide sequence is entirely transformed, with the plDDT for the 39-residue design increasing from 53 to 95. Peptides ranging from 25 to 40 residues in length were designed in as little as 24 h, all displaying high plDDT scores (Figure 1b) and strong agreement with adversarial validation using AlphaFold-Multimer [18] (Methods; Figure 1c, e).

A top candidate of 39 residues in length (Figure 1a, d) was synthesised and assessed in cellular assays for its ability to inhibit HIV-1 infection by the two reference strains NL4-3 and AD8 (Methods). The designed cyclic peptide engages the hexameric form of gp41, a conformation that only becomes transiently accessible at the precise moment of membrane fusion (Figure 1a). This brief exposure limits the opportunity for inhibition, meaning that even potent binders may exhibit reduced apparent efficacy in cellular assays. Nonetheless, the observed inhibition indicates successful target engagement under these constrained conditions, with low micromolar IC_50_ values recorded for both HIV-1 strains (6.18 μM for AD8 and 9.40 μM for NL4-3; Figure 1f).

Importantly, no cellular toxicity was observed, and cell viability remained high even at elevated peptide concentrations across 24, 48, and 72 hours post-exposure (Figure 1g). This indicates that the peptide acts selectively on the virus without harming host cells. These findings underscore the potential of cyclic peptides to exploit brief but critical windows of vulnerability in the viral life cycle, while avoiding off-target effects. The integrated workflow - combining rapid AI-driven design with experimental validation - demonstrates a promising approach for the swift development of peptide-based therapeutics capable of keeping pace with the virus’s continual evolution.

### TDF as a Reference for Antiviral Activity

As a positive control, we used Tenofovir Disoproxil Fumarate (TDF) [37], a clinically approved reverse transcriptase inhibitor that acts post-entry by blocking viral replication [38]. In contrast, the designed cyclic peptide intervenes at an earlier stage of the HIV life cycle by preventing membrane fusion through binding to gp41. Although TDF exhibited lower IC_50_ values, this is expected due to its broader temporal window of activity during viral replication. The comparatively higher IC_50_ of the cyclic peptide (Figure 2a) reflects the narrow window during which gp41 adopts its fusion-active conformation, thereby limiting opportunities for inhibitor binding. Nevertheless, the observed efficacy of the peptide confirms successful target engagement during this transient window and supports its potential role in a multi-pronged antiviral strategy, particularly in cases of resistance to reverse transcriptase inhibitors.

**Figure 2.**
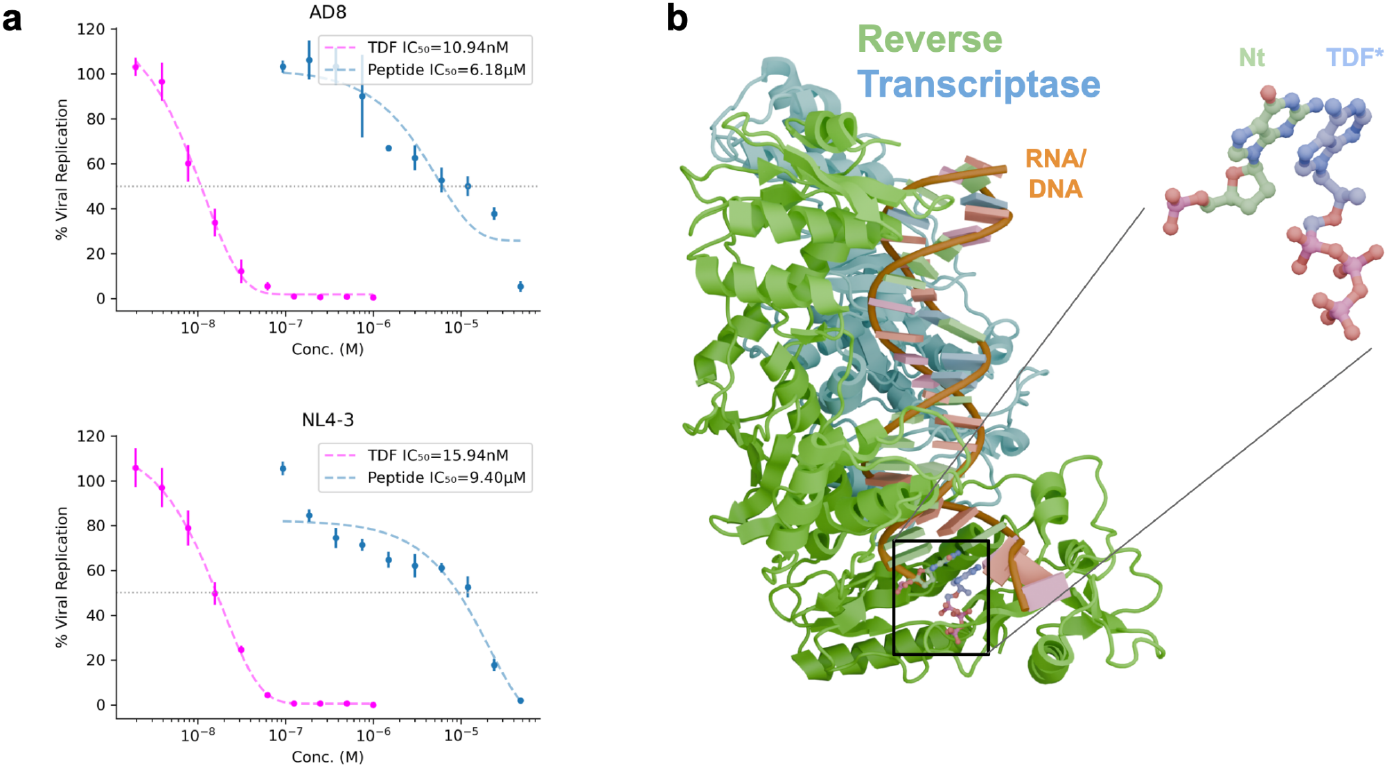
Comparison with the Reverse Transcriptase (RT) inhibitor Tenofovir Disoproxil Fumarate (TDF). **a)** Dose-response curves for the designed cyclic peptide (L39) and the positive control TDF. Curves are colour-coded to distinguish between compounds with median (points) and standard deviations (vertical lines) at each measured concentration, based on n=4 measurements per concentration. The fitted IC_50_ values for TDF were 11 nM against AD8 and 16 nM against NL4-3 HIV-1 strains. The designed cyclic peptide (L39) show IC_50_ values of 6.18 μM (AD8) and 9.40 μM (NL4-3) under the same conditions. The point of 50% viral replication is marked with a dotted line. While TDF exhibit lower IC_50_ values, this is expected given the broader temporal window of activity during the viral replication cycle. The comparatively higher IC_50_ of the cyclic peptide reflects the narrow window during which gp41 adopts its fusion-active conformation, thereby limiting the opportunity for inhibitor binding. **b)** Structure of a TDF analogue (TDF*, tenofovir diphosphate) in complex with HIV-1 reverse transcriptase (PDB ID: 1T05). TDF mimics a natural nucleotide (Nt) and binds at the enzyme’s active site, blocking the incorporation of subsequent nucleotides and thereby halting viral DNA synthesis and replication.

TDF engages reverse transcriptase by mimicking natural nucleotides (Figure 2b), thereby terminating viral DNA synthesis after the virus has entered the host cell. Its mechanism of action is well established and benefits from extensive structural characterisation developed over decades [38–41]. In contrast, our approach introduces an entirely new mode of inhibition by designing a cyclic peptide that binds to the gp41 fusion hexamer - an interface not previously targeted. Remarkably, this was achieved without any prior structural data, relying solely on the amino acid sequence of the target protein. This demonstrates the power of AI-driven de novo design to uncover novel mechanisms of action and develop inhibitors against conserved yet structurally elusive targets, offering a valuable complement to existing antiretroviral therapies.

### Mutational Impact and Structural Comparison to Known Binders

To address the challenge of drug resistance, the designed cyclic peptide specifically avoids the enfuvirtide binding site on gp41, which is prone to mutations such as N43D [33]. We based our design on the sequence from PDB ID 3CP1, which includes this resistance mutation and reveals the resulting structural alterations in the six-helix bundle. Importantly, although the N43D mutation interferes with local hydrogen bonding and leads to a slight unwinding of the HR2 helix, the rest of the helical core retains its overall structure, showing an RMSD of just 0.81 Å compared to the wild type [33]. EvoBind selected a binding site on a separate, structurally conserved surface distant from the mutation (Figure 3a). This enables the cyclic peptide to target a region of gp41 that retains its structural integrity even in the presence of common mutations, offering a more robust and durable mode of inhibition.

**Figure 3.**
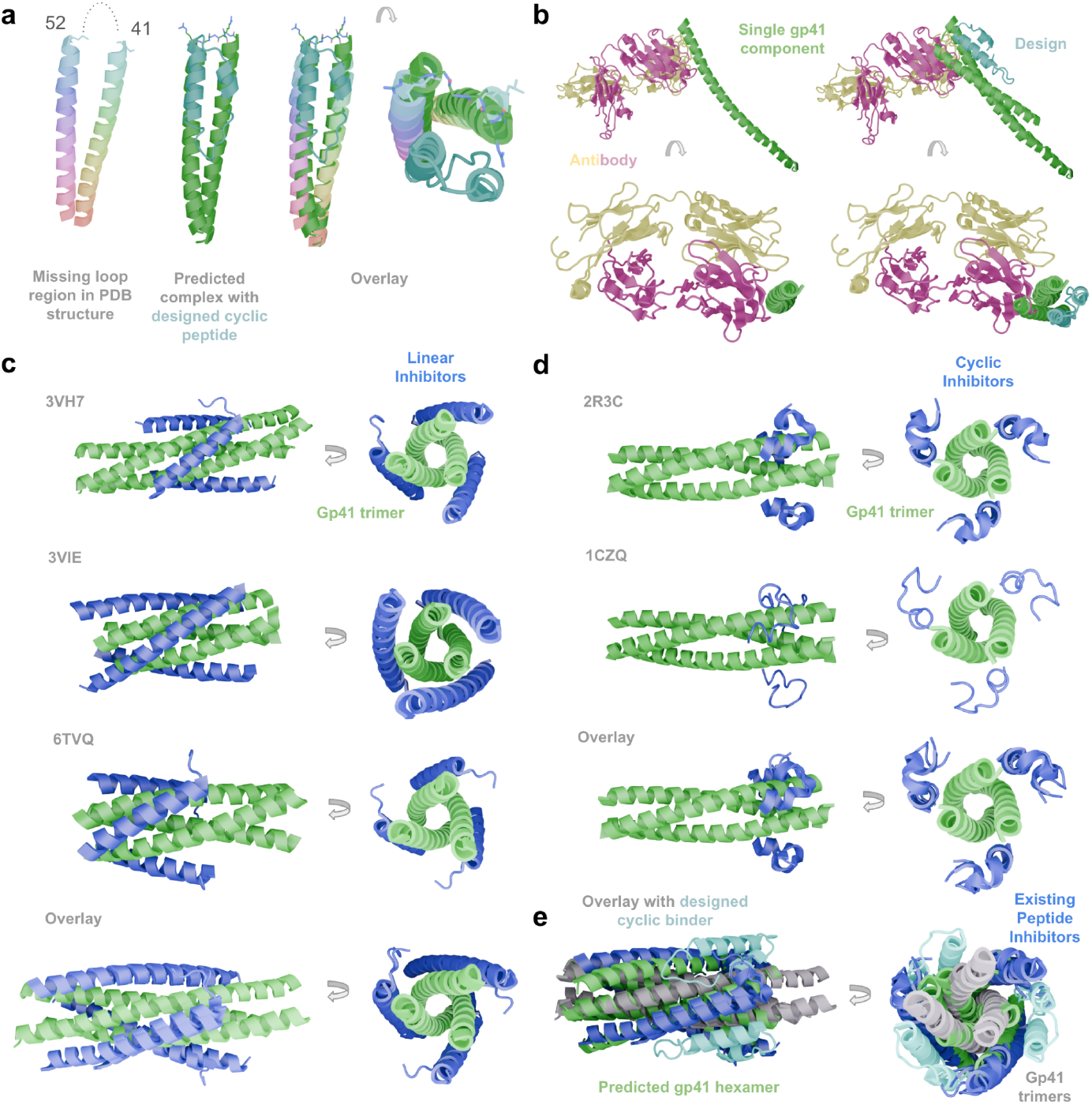
A novel binding mode that avoids resistance mutations. **a)** The resistance mutation N43D, associated with the drug Enfuvirtide, is located in a flexible loop region of gp41 that is unresolved in the crystal structure (PDB ID 3CP1), shown here in a rainbow gradient. In contrast, the predicted complex with the designed cyclic peptide of length 39 includes this loop but reveals that it does not contribute to the predicted peptide interface, suggesting that the design is not affected by this resistance-associated region. **b)** Structure of single gp41 helix component with an inhibitory antibody (PDB ID 4KHX). The antibody targets the individual gp41 component, but on the opposite side of the designed cyclic peptide which targets a novel interface created by two gp41 components according to the predicted structure. **c)** There are many gp41 peptide inhibitors with solved structure in the PDB (displayed are structures from PDB IDs 3VH7, 3VIE and 6TVQ) that utilise components of the gp41 hexamer, specifically the outer components (C-HR or HR2). All of these thereby inhibit hexamer formation in the same way, distinct from the designed cyclic peptide which targets the gp41 hexamer as a complex (e). **d)** Similar to the linear peptide inhibitors, cyclic inhibitors exist that also target the inner gp41 trimeric component N-HR or HR1 (PDB IDs 2R3C and 1CZQ, both highly similar). **e)** Overlay showing the novelty of our design: the cyclic peptide binder generated by EvoBind (light blue) targets a unique interface formed by two gp41 helices (HR1 and HR2 in green), distinct from the binding sites (grey) exploited by previously described inhibitors (blue).

Compared to other known binders targeting gp41, the designed cyclic peptide targets a different interface with a novel binding mode. There are antibodies (Figure 3b) that bind to the single components of the gp41 hexamer, and several peptide analogues derived from gp41 itself that target the trimeric state by mimicking the hexamer (Figure 3c). Other peptides are cyclic and target the same interface created by the inner three gp41 components (Figure 3d). However, none of these binders target the fusion hexamer, highlighting the possibility of EvoBind to target gp41 in a novel way with a cyclic peptide that is not simply mimicking the structural components of gp41 (Figure 3e).

## Discussion

The rapid emergence of drug resistance in HIV-1 continues to undermine the long-term effectiveness of current therapies. As mutations accumulate in key viral proteins, many standard inhibitors lose potency, underscoring the need for continuous innovation in antiviral drug development. To meet this challenge, we use EvoBind, an artificial intelligence-driven design framework that enables the de novo generation of peptide therapeutics directly from protein sequence information. By bypassing the need for experimentally determined structures or known binding sites, EvoBind opens new avenues for targeting conserved regions of viral proteins. This capability is particularly valuable in settings where rapid adaptation is essential, offering a scalable strategy for addressing resistance and accelerating early-stage therapeutic discovery.

Cyclic peptides, in particular, are promising drug candidates due to their favourable pharmacological properties, including improved proteolytic stability and the potential for oral bioavailability. Their conformational rigidity and ability to engage protein-protein interfaces make them especially well suited to targeting viral proteins involved in entry and fusion. Here, we demonstrate the application of EvoBind [23] for the rapid generation of cyclic peptide binders targeting the HIV-1 fusion protein gp41. The resulting design represents an entirely new way of targeting gp41 according to the predicted structure (Figure 3).

Starting with only the target amino acid sequence, we generated candidate binders in a single design round, which demonstrated clear antiviral activity in cells. The resulting IC_50_ values were low micromolar, 6.18 and 9.40 μM for HIV strains AD8 and NL4-3, respectively, consistent with the transient exposure of the hexameric gp41 core structure. Importantly, no cytotoxicity was observed in treated cells. While the measurements do not follow a strict sigmoidal shape, we do not believe this impacts the IC_50_ values, as the fitted curves closely align with the measured data points within the IC_50_ region, corresponding to the linear portion of the sigmoidal response. Had higher concentrations been included in the assay, the full sigmoidal curve would have emerged, as seen for TDF (Figure 2).

The success of EvoBind, which relies solely on target sequence information, suggests that the approach can be generalised to other viral pathogens. By enabling rapid, single-round generation of binders, EvoBind offers a scalable and adaptive strategy to address both emerging resistance and future pandemic threats. This work highlights the potential of modern de novo binder design technologies to accelerate early-stage drug discovery. When combined with streamlined experimental testing, artificial intelligence-guided design can reduce development timelines, lower costs, and deliver timely therapeutic solutions in response to viral evolution and global health challenges.

## Methods

### Cyclic peptide design with EvoBind towards gp41

EvoBind (version 2) was obtained from GitHub (https://github.com/patrickbryant1/EvoBind) and run using the default settings, which include eight recycles and multiple sequence alignment (MSA) generation via HHblits [42] against the uniclust30_2018_08 database [43]. All computations were performed on a single NVIDIA A100 Tensor Core GPU with 40 GB of RAM. As input, we used the full FASTA sequence of chain A from the gp41 structure (PDB ID: 3CP1, see “gp41 sequence”). This sequence represents a fusion of two peptide segments from the gp41 hexamer, thereby forming a “pseudo-dimer” suitable for binder design. Figure 4 provides a visual explanation of the design procedure.

**Figure 4.**
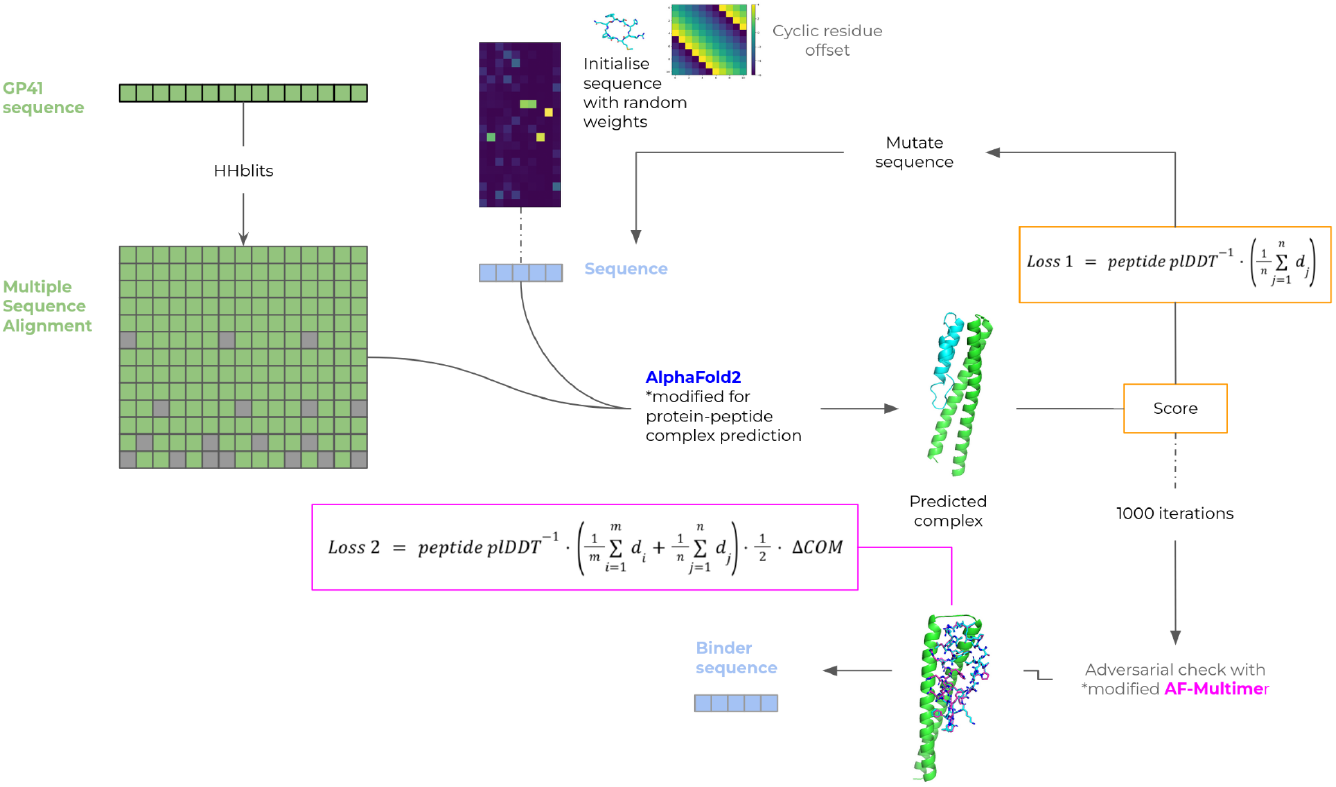
Design procedure for peptide binders using EvoBind2. The only input to the design pipeline is the target protein sequence for gp41; no prior information about the binding site or the peptide length is required. The target sequence is searched against the Uniclust30 database using HHblits to generate a multiple sequence alignment (MSA). The peptide sequence is randomly initialised by sampling weights between 0 and 1 from a Gumbel distribution, with the argmax at each position (L×20) determining the initial amino acid sequence. We also use a cyclic offset to inform the structure prediction that a cyclic peptide is being designed. This initial peptide and the MSA are then input into a modified version of AlphaFold2 to predict the structure of the protein-peptide complex. A scoring function (Loss 1) is applied to the prediction. If the score improves upon the previous best, the current peptide is retained and used as the basis for introducing a new mutation, followed by another round of structure prediction. This process is repeated for 1000 mutation cycles. The resulting AlphaFold2 predictions are then compared with those generated by a modified AlphaFold-Multimer model (Loss 2) to ensure agreement, and the best candidate is experimentally validated.

We explicitly aimed to design cyclic peptides ranging from 25 to 40 amino acids in length. This range was chosen based on the precedent set by T20 (Enfuvirtide), a clinically approved 36-residue linear peptide that targets gp41 [44], supporting the hypothesis that a length in this range is suitable for effective inhibition. EvoBind was run with five distinct random initialisations for each of the sixteen lengths, for a total of 1000 design iterations (n=5·16·1000=80000). The design process was conducted in an untargeted manner, addressing the entire gp41 target sequence and each run finished within 24 h. Figure 5 displays the optimisation procedure for each length.

**Figure 5.**
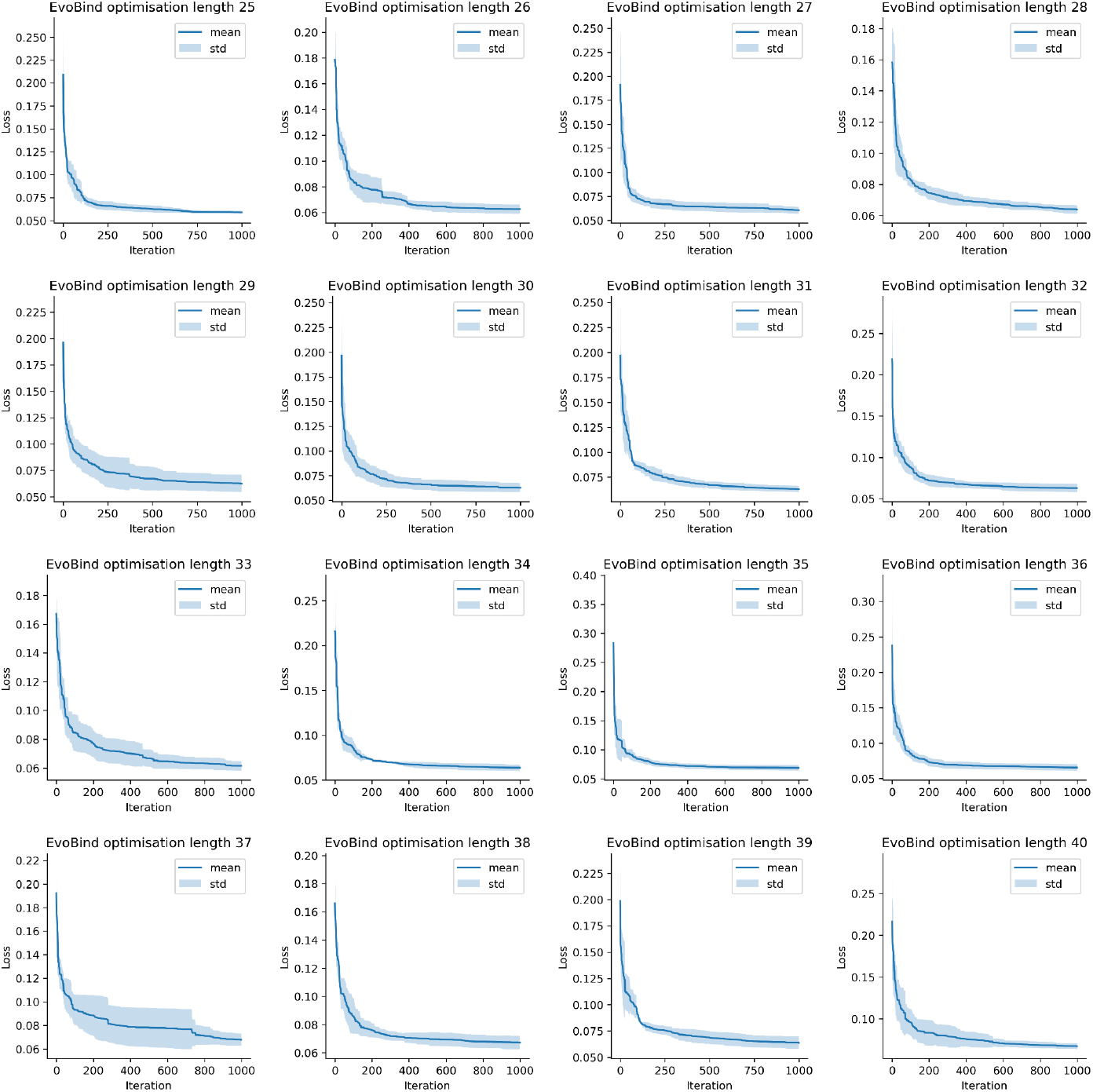
Best loss vs iteration. Lowest loss for the design (equation 1) so far vs iteration for the design of different lengths. The average across five different initialisations is displayed as a line with shaded standard deviation.

#### gp41 sequence, 86 amino acids

TLTVQARQLLSGIVQQQNDLLRAIEAQQHLLQLTVWGIKQLQARSGGRGGWMEWDREINNYTS LIHSLIEESQNQQEKNEQELLEL

The loss function used for the optimisation is defined as:

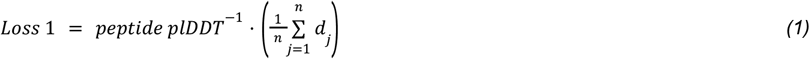

Here, the peptide plDDT (the predicted local distance difference test [45]) refers to the average plDDT score across the entire peptide, while dj denotes the shortest distance between any atom nnn in the peptide and any atom within the target protein. The plDDT score reflects the local reliability of predicted protein structures, with higher values corresponding to greater confidence in the accuracy of the model.

The loss for the adversarial selection is defined as:

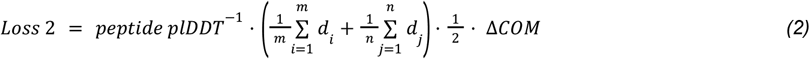

Here, the peptide plDDT refers to the average predicted local distance difference test score across the peptide. di represents the shortest distance between any atom in the receptor target atoms (defined as Cβ atoms within 8 Å of the peptide, as predicted by EvoBind2) and any atom in the peptide. dj is the shortest distance between any atom in the peptide and any atom in the receptor target atoms. ΔCOM denotes the distance between the alpha carbon centres of mass of the predicted peptides from the design and validation procedures, respectively.

### Binder design selection

From the output of EvoBind, we selected the top 10% of peptide models with the lowest design loss *(equation 1)* and a plDDT score above 90 (n=7447). These top candidates were further validated in silico using AlphaFold-Multimer in an adversarial manner. The rationale is that it is unlikely for two distinct models, trained on separate data sources, to produce high-confidence predictions for an incorrect structure. Additional filters were then applied using the adversarial predictions: the top 10% of candidates with the lowest combined loss *(equation 2)* and a plDDT score of at least 90 in both the EvoBind and AlphaFold-Multimer predictions (for the same peptide sequence) were retained. We selected the top 2 design candidates for synthesis (Table 1). However, only the second-best design of length 39 could easily be synthesised by our supplier, Genscript, at high purity (>90%). All analyses were therefore conducted using only this cyclic peptide.

**Table 1.** Metrics for the top two selected designs. Only length 39 could be easily synthesised at high purity. AFM loss corresponds to equation 2, and Evo loss to equation 1 and the combined loss is simply the sum of the two.

| Length | AFM pIDDT | AFM loss | Evo pIDDT | Evo loss | Sequence | Combined loss | Purity (%) |
| --- | --- | --- | --- | --- | --- | --- | --- |
| 38 | 93.8 | 0.00471 | 92.2 | 0.05927 | GMDVPWTAMKYDAGFEGGS<br>YMGHAQSQVAETSARLSAW | 0.06398 | NA |
| 39 | 96.1 | 0.00270 | 95.3 | 0.06200 | MTPLVNAV PADNGDAVLLKR<br>CWAHYKDNCFGFPLYKPVD | 0.06470 | 90.4 |

### Net peptide content determination with UV spectroscopy

The peptide was lyophilised with TFA salt and water, leading to a lower content than the 90.4% purity before lyophilisation. To analyse the actual peptide content, we diluted the lyophilised powder to 1 mM and used UV spectroscopy with Nanodrop. The designed sequence contains the following amino acids that absorb light at 280 nM:

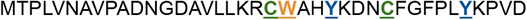

The extinction coefficients [46] are calculated accordingly:

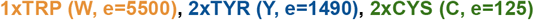

Resulting in a combined extinction coefficient of:

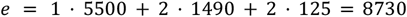

We measured the absorbance twice, obtaining absorbance values of 4.20 and 4.12 with an average of 4.16. The absorbance is related to the concentration (c) and path length (l=1 mm) in the following way:

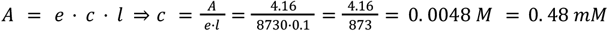

This means that all concentrations prepared from diluting the lyophilised powder actually contain 48% peptide and should be adjusted accordingly.

### Compounds

The designed cyclic peptide was synthesised with >90% purity (verified by HPLC and mass spectrometry, Supplementary Material). Tenofovir disoproxil fumarate (TDF, Selleckchem, S1400), a nucleotide reverse transcriptase inhibitor that blocks HIV-1 replication by inhibiting viral DNA synthesis, was used as a reference compound. Both compounds were initially dissolved in DMSO, followed by dilution in complete cell culture medium to the required concentrations for assays, with a final DMSO concentration not exceeding 0.5%.

### Cells and viruses

The TZM-bl and HEK-293T cell lines (NIH AIDS Research and Reference Reagent Program, USA) were cultured in DMEM (Sigma, USA) supplemented with 10% FBS, penicillin/streptomycin (100 IU/mL and 50 mg/mL, respectively), and 2 mM L-glutamine. HIV-1 reference strains NL4-3 (X4-tropic, T-cell line adapted laboratory strain) and AD8 (R5-tropic, macrophage-tropic primary isolate) were obtained through the NIH AIDS Reagent Program. Viral stocks of HIV-1_AD8_ and HIV-1_NL4.3_ laboratory strains were obtained by transient transfection of their corresponding plasmids in HEK-293T cells (ATCC, Manassas, VA, USA). Supernatants were collected at 48 h and 72 h. Viral stocks were clarified by centrifugation before evaluating the viral titer by HIV-1 p24gag ELISA kit (INNOTEST® Innogenetics, Belgium). Viral infectivity was assessed by tissue culture infective doses (TCID50) using the Spearman-Kärber method based on the results from the HIV-1 p24gag ELISA kit.

### Cytotoxicity assay

Cytotoxicity was determined using the CellTiter-Glo 2.0 Luminescent Cell Viability Assay (Promega, Madison, WI, USA) according to the manufacturer’s protocol. Briefly, TZM-bl cells were seeded at a density of 10,000 cells per well in 96-well plates and allowed to adhere for 24 hours prior to treatment. Compounds were prepared in a 10-fold serial dilution series in complete medium, with concentrations ranging from 48 μM. Negative control wells containing an equivalent concentration of DMSO were included on each plate to establish baseline cell death. At 24, 48, and 72 hours post-treatment, CellTiter-Glo reagent (equal volume to the culture medium) was added to each well. The luminescent signal, proportional to the amount of cellular ATP reflecting viable cell numbers, was measured. Luminescence measurements were background-corrected, and relative cell viability was calculated as a percentage. Dose-response curves and the half-maximal cytotoxic concentration (CC_50_) were calculated using GraphPad Prism software (version 9.0). The designed cyclic peptide of length 39 was tested in duplicate in each experiment. For subsequent antiviral assays, the highest non-cytotoxic concentration (defined as the concentration yielding 90–100% cell viability) was established as the maximum test concentration.

### HIV-1 antiviral assays

The antiviral activity of the designed cyclic peptide of length 39 was evaluated against HIV-1 reference strains NL4-3 and AD8 using a TZM-bl cell-based phenotypic assay. TZM-bl cells (10,000 cells/well) were infected with wild-type strains at a 150 tissue culture infective dose (TCID50) in the presence of 2-fold serial dilutions of the peptide (48 μM to 0.1875 μM). Each assay plate included tenofovir (TDF, Sellckchem, USA) as a reference compound, uninfected cell controls, and virus-only controls. After 48 hours of incubation, cells were lysed and viral infection was quantified by measuring luciferase expression using the Bright-Glo Luciferase Assay system (Promega, USA). Luminescence signals were normalised to virus-only controls (100% infection) and uninfected controls (0% infection). All assays were performed in duplicate and validated across two independent experiments.

### IC_50_ calculations

Dose-response curves were fitted using a sigmoidal function of the form:

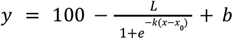

The resulting fits were then used to calculate the half-maximal inhibitory concentration (IC_50_) values shown in Figures 1 and 2. Although the measurements for the designed peptide is not perfectly sigmoidal, we do not believe this affects the IC_50_ values, as the fitted curves accurately follow the measured data points in the IC_50_ region.

For the NL4-3 strain treated with the designed peptide, the IC_50_ was approximately 9.4 μM, based on fitting parameters L=105.55, x_0_=2.25 ·10^-6^, k=1, and b=1.86. For the AD8 strain, the IC_50_ was approximately 6.2 μM, with fitting parameters L=106.13, x_0_=2.25?·10^-6^, k=1, and b=5.43.

## Data Availability

All raw data are available at https://zenodo.org/uploads/15234340

## Code Availability

The code used for analysis and figure generation can be found at: https://gitlab.com/patrickbryant1/gp41

The software used for peptide binder design (EvoBind) can be found at: https://github.com/patrickbryant1/EvoBind

## Contributions

D.D., Q.L. and P.B. conceived and designed the study. D.D. and P.B. drafted the initial manuscript. Binder design was performed by D.D. in coordination with Q.L. and P.B. HIV inhibition and toxicity assays were conducted by F.G. and R.D. All authors contributed to the analysis and the preparation of the final manuscript. P.B. and A.S. obtained funding.

## Acknowledgements

This study was supported by the SciLifeLab & Wallenberg Data Driven Life Science Program (grant: KAW 2020.0239, P.B). Computational resources were enabled by the supercomputing resource Berzelius provided by National Supercomputer Centre at Linköping University and the Knut and Alice Wallenberg foundation with project ids Berzelius-2023-267, Berzelius-2024-78, Berzelius-2024-292 and berzelius-2025-41 (P.B). This project has received funding through Karolinska Institutet Research Foundation (2024-02416) and Stiftelsen Läkare Mot AIDS Forskningsfond (FOa2023-0019) to RCD and the Swedish Research Council (2020–02129) and Stockholm County Council (FoUI-955284 and FoUI-953887) to A.S.

## Supplementary information

**Supplementary Figure 1.**
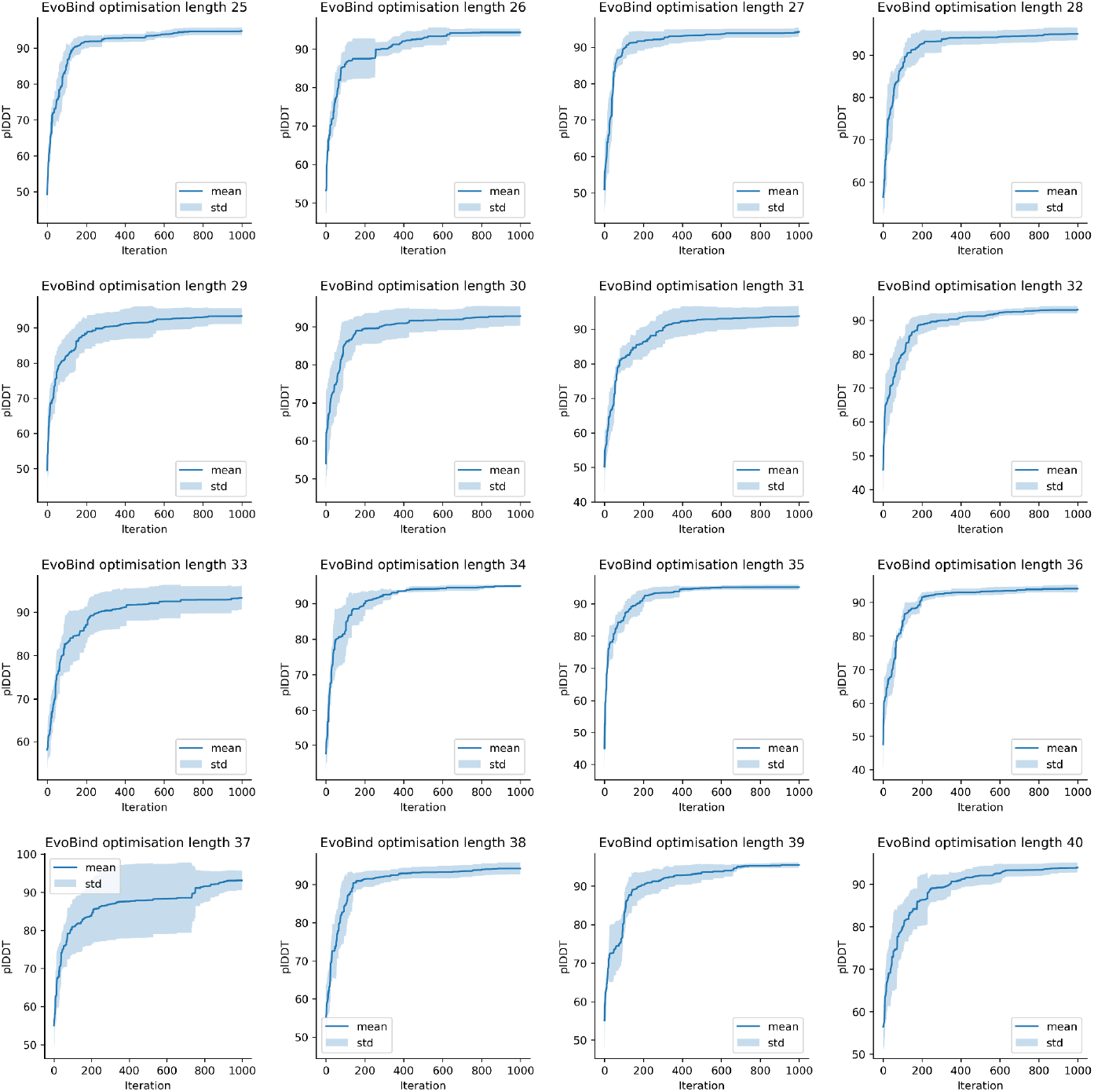
Highest plDDT vs iteration. Highest predicted confidence of the peptide (plDDT) so far vs iteration for the design of different lengths. The average across five different initialisations is displayed as a line with shaded standard deviation.

## References

1. Global HIV & AIDS statistics — Fact sheet. [cited 16 Apr 2025]. Available: https://www.unaids.org/en/resources/fact-sheet

2. HIV data and statistics. [cited 16 Apr 2025]. Available: https://www.who.int/teams/global-hiv-hepatitis-and-stis-programmes/hiv/strategic-information/hiv-data-and-statistics

3. GBD 2021 HIV Collaborators. Global, regional, and national burden of HIV/AIDS, 1990-2021, and forecasts to 2050, for 204 countries and territories: the Global Burden of Disease Study 2021. Lancet HIV. 2024;11: e807–e822.

4. Van Duyne R, Kuo LS, Pham P, Fujii K, Freed EO. Mutations in the HIV-1 envelope glycoprotein can broadly rescue blocks at multiple steps in the virus replication cycle. Proceedings of the National Academy of Sciences of the United States of America. 2019;116: 9040.

5. Beretta M, Migraine J, Moreau A, Essat A, Goujard C, Chaix M-L, et al. Common evolutionary features of the envelope glycoprotein of HIV-1 in patients belonging to a transmission chain. Scientific Reports. 2020;10: 1–17.

6. Aquaro S, D’Arrigo R, Svicher V, Perri GD, Caputo SL, Visco-Comandini U, et al. Specific mutations in HIV-1 gp41 are associated with immunological success in HIV-1-infected patients receiving enfuvirtide treatment. J Antimicrob Chemother. 2006;58: 714–722.

7. Rossi E, Meuser ME, Cunanan CJ, Cocklin S. Structure, Function, and Interactions of the HIV-1 Capsid Protein. Life. 2021;11: 100.

8. Rihn SJ, Wilson SJ, Loman NJ, Alim M, Bakker SE, Bhella D, et al. Extreme Genetic Fragility of the HIV-1 Capsid. PLoS Pathogens. 2013;9: e1003461.

9. Baeten JM. Lenacapavir for Human Immunodeficiency Virus (HIV) Prevention: A Commitment to Equitable Access and Partnership by Gilead Sciences. Clin Infect Dis. 2025. doi:10.1093/cid/ciaf116

10. Kelley CF, Acevedo-Quiñones M, Agwu AL, Avihingsanon A, Benson P, Blumenthal J, et al. Twice-Yearly Lenacapavir for HIV Prevention in Men and Gender-Diverse Persons. New England Journal of Medicine. 2025 [cited 16 Apr 2025]. doi:10.1056/NEJMoa2411858

11. Bester SM, Adu-Ampratwum D, Annamalai AS, Wei G, Briganti L, Murphy BC, et al. Structural and Mechanistic Bases of Viral Resistance to HIV-1 Capsid Inhibitor Lenacapavir. mBio. 2022 [cited 28 Apr 2025]. doi:10.1128/mbio.01804-22

12. Margot N, Vanderveen L, Naik V, Ram R, Parvangada PC, Martin R, et al. Phenotypic resistance to lenacapavir and monotherapy efficacy in a proof-of-concept clinical study. The Journal of antimicrobial chemotherapy. 2022;77. doi:10.1093/jac/dkab503

13. Margot NA, Jogiraju V, Pennetzdorfer N, Naik V, VanderVeen LA, Ling J, et al. Resistance Analyses in Heavily Treatment-Experienced People with HIV Treated with the Novel HIV Capsid Inhibitor Lenacapavir After 2 years. The Journal of infectious diseases. 2025 [cited 28 Apr 2025]. doi:10.1093/infdis/jiaf050

14. Jumper J, Evans R, Pritzel A, Green T, Figurnov M, Ronneberger O, et al. Highly accurate protein structure prediction with AlphaFold. Nature. 2021;596: 583–589.

15. Bryant P, Noé F. Structure prediction of alternative protein conformations. Nature Communications. 2024;15: 1–12.

16. Lewis S, Hempel T, Jiménez-Luna J, Gastegger M, Xie Y, Foong AYK, et al. Scalable emulation of protein equilibrium ensembles with generative deep learning. bioRxiv. 2025. p. 2024.12.05.626885. doi:10.1101/2024.12.05.626885

17. Bryant P, Pozzati G, Elofsson A. Improved prediction of protein-protein interactions using AlphaFold2. Nature Communications. 2022;13: 1–11.

18. Evans R, O’Neill M, Pritzel A, Antropova N, Senior A, Green T, et al. Protein complex prediction with AlphaFold-Multimer. bioRxiv. 2022. p. 2021.10.04.463034. doi:10.1101/2021.10.04.463034

19. Abramson J, Adler J, Dunger J, Evans R, Green T, Pritzel A, et al. Accurate structure prediction of biomolecular interactions with AlphaFold 3. Nature. 2024;630: 493–500.

20. Bryant P, Kelkar A, Guljas A, Clementi C, Noé F. Structure prediction of protein-ligand complexes from sequence information with Umol. Nature Communications. 2024;15: 1–12.

21. Krishna R, Wang J, Ahern W, Sturmfels P, Venkatesh P, Kalvet I, et al. Generalized biomolecular modeling and design with RoseTTAFold All-Atom. Science. 2024 [cited 16 Apr 2025]. doi:10.1126/science.adl2528

22. Qiao Z, Nie W, Vahdat A, Miller TF, Anandkumar A. State-specific protein–ligand complex structure prediction with a multiscale deep generative model. Nature Machine Intelligence. 2024;6: 195–208.

23. Li Q, Vlachos EN, Bryant P. Design of linear and cyclic peptide binders of different lengths from protein sequence information. bioRxiv. 2024. p. 2024.06.20.599739. doi:10.1101/2024.06.20.599739

24. Pacesa M, Nickel L, Schellhaas C, Schmidt J, Pyatova E, Kissling L, et al. BindCraft: one-shot design of functional protein binders. bioRxiv. 2024. p. 2024.09.30.615802. doi:10.1101/2024.09.30.615802

25. Sappington I, Toul M, Lee DS, Robinson SA, Goreshnik I, McCurdy C, et al. Improved protein binder design using beta-pairing targeted RFdiffusion. bioRxiv. 2024. p. 2024.10.11.617496. doi:10.1101/2024.10.11.617496

26. Callaway E. AlphaFold is running out of data — so drug firms are building their own version. In: Nature Publishing Group UK [Internet]. 27 Mar 2025 [cited 16 Apr 2025]. doi:10.1038/d41586-025-00868-9

27. Škrinjar P, Eberhardt J, Durairaj J, Schwede T. Have protein-ligand co-folding methods moved beyond memorisation? bioRxiv. 2025. p. 2025.02.03.636309. doi:10.1101/2025.02.03.636309

28. Muttenthaler M, King GF, Adams DJ, Alewood PF. Trends in peptide drug discovery. Nature Reviews Drug Discovery. 2021;20: 309–325.

29. Wang L, Wang N, Zhang W, Cheng X, Yan Z, Shao G, et al. Therapeutic peptides: current applications and future directions. Signal Transduction and Targeted Therapy. 2022;7: 1–27.

30. Merz ML, Habeshian S, Li B, David J-AGL, Nielsen AL, Ji X, et al. De novo development of small cyclic peptides that are orally bioavailable. Nature Chemical Biology. 2023;20: 624–633.

31. Liu L, Yang L, Cao S, Gao Z, Yang B, Zhang G, et al. CyclicPepedia: a knowledge base of natural and synthetic cyclic peptides. Brief Bioinform. 2024;25. doi:10.1093/bib/bbae190

32. Brown DG, Wobst HJ, Kapoor A, Kenna LA, Southall NT. Clinical development times for innovative drugs. Nature reviews Drug discovery. 2022;21: 793.

33. Bai X, Wilson KL, Seedorff JE, Ahrens D, Green J, Davison DK, et al. Impact of the Enfuvirtide Resistance Mutation N43D and the Associated Baseline Polymorphism E137K on Peptide Sensitivity and Six-Helix Bundle Structure†‡. 2008 [cited 16 Apr 2025]. doi:10.1021/bi702509d

34. Taylor BS, Hunt G, Abrams EJ, Coovadia A, Meyers T, Sherman G, et al. Rapid Development of Antiretroviral Drug Resistance Mutations in HIV-Infected Children Less Than Two Years of Age Initiating Protease Inhibitor-Based Therapy in South Africa. AIDS Research and Human Retroviruses. 2011;27: 945.

35. Doms RW, Moore JP. HIV-1 Membrane Fusion: Targets of Opportunity. The Journal of Cell Biology. 2000;151: f9.

36. Viral surface glycoproteins, gp120 and gp41, as potential drug targets against HIV-1: Brief overview one quarter of a century past the approval of zidovudine, the first anti-retroviral drug. European Journal of Medicinal Chemistry. 2011;46: 979–992.

37. Tenofovir disoproxil. Wikimedia Foundation, Inc.; 14 Dec 2004 [cited 23 Apr 2025]. Available: https://en.wikipedia.org/wiki/Tenofovir_disoproxil

38. Tuske S, Sarafianos SG, Clark AD, Ding J, Naeger LK, White KL, et al. Structures of HIV-1 RT–DNA complexes before and after incorporation of the anti-AIDS drug tenofovir. Nature Structural & Molecular Biology. 2004;11: 469–474.

39. Gu W, Martinez S, Nguyen H, Xu H, Herdewijn P, De Jonghe S, et al. Tenofovir-Amino Acid Conjugates Act as Polymerase Substrates—Implications for Avoiding Cellular Phosphorylation in the Discovery of Nucleotide Analogues. Journal of Medicinal Chemistry. 2020 [cited 23 Apr 2025]. doi:10.1021/acs.jmedchem.0c01747

40. The structure of HIV-1 reverse transcriptase complexed with 9-chloro-TIBO: lessons for inhibitor design. Structure. 1995;3: 915–926.

41. Structure of HIV-1 reverse transcriptase in a complex with the non-nucleoside inhibitor α-APA R 95845 at 2.8 Å resolution. Structure. 1995;3: 365–379.

42. Steinegger M, Meier M, Mirdita M, Vöhringer H, Haunsberger SJ, Söding J. HH-suite3 for fast remote homology detection and deep protein annotation. BMC Bioinformatics. 2019;20: 473.

43. Mirdita M, von den Driesch L, Galiez C, Martin MJ, Söding J, Steinegger M. Uniclust databases of clustered and deeply annotated protein sequences and alignments. Nucleic Acids Res. 2017;45: D170–D176.

44. Champagne K, Shishido A, Root MJ. Interactions of HIV-1 Inhibitory Peptide T20 with the gp41 N-HR Coiled Coil. The Journal of Biological Chemistry. 2009;284: 3619.

45. Mariani V, Biasini M, Barbato A, Schwede T. lDDT: a local superposition-free score for comparing protein structures and models using distance difference tests. Bioinformatics. 2013;29: 2722.

46. Thermo Fisher extinction coefficients. [cited 25 Apr 2025]. Available: https://assets.thermofisher.com/TFS-Assets/LSG/Application-Notes/TR0006-Extinction-coefficients.pdf

